# Self-paced treadmill controller algorithm based on position and speed of centre of mass

**DOI:** 10.1101/2022.06.21.496740

**Authors:** Hossein Mokhtarzzadeh, Rosie Richards, Thomas Geijtenbeek

## Abstract

**Background:** Self-paced treadmills are increasingly used in clinical and research settings. Using self-paced (SP) treadmills, researchers can simulate overground walking while participants can walk with different but comfortable gait speeds in a controlled environment. Several algorithms have been designed for self-paced treadmills based on data from force plates, motion capture, and even marker-less systems such as 3D depth cameras.

**Methods:** We present a non-linear controller that implements a self-paced algorithm integrated with treadmills. This algorithm uses the subject’s centre-of-mass (CoM) position and velocity, relative to the front and back end of the treadmill as inputs. The controller continuously adjusts the treadmill’s belt speed via belt acceleration. The algorithm attempts to prevent the subject reaching the front and back of treadmill via minimal treadmill acceleration.

**Findings:** This method has been safely used in previous studies with over 410 subjects in various populations. We simulated the use of the SP algorithm with three different sensitivities (0.2, 1 and 2). The belt speed predicted by algorithm simulation in matched well with the belt speeds of experiments in (Gait Realtime Analysis Interactive Lab (GRAIL) system.

**Interpretation:** This algorithm is integrated with a VR environment in which the subject can be immersed and even be mechanically perturbed. Additionally, this algorithm can be implemented in other treadmills where CoM position is known. We encourage researchers to use and build upon our well-established SP algorithm toward a more standardized SP algorithm in different gait scenarios across various instrumented treadmills with different populations.

## Introduction

Self-paced (SP) treadmills can play a key role in the assessment of gait performance in the laboratory in different populations (Bahadori et al., 2021; Canete and Jacobs, 2021; Castano and Huang, 2021; Theunissen et al., 2022b; van der Krogt et al., 2014; Wei et al., 2020). Self-paced experiments may offer benefits over field based or fixed-speed treadmill studies. Field studies are time-consuming and result in a less controlled environment. Fixed-speed treadmill studies may cause compensatory mechanisms to keep the fixed speed which could affect gait parameters (Sinitski et al., 2015). Gait patterns in SP and fixed-speed treadmill studies were comparable in terms of kinematics, kinetics, and energetics across gait speeds (Sloot et al., 2014; Theunissen et al., 2022b); however, when virtual reality (VR) visual flow was present, subjects could reach a steady state of gait speed more rapidly in SP experiments (Plotnik et al., 2015). Additionally, most studies on SP treadmills have been on a small population of usually healthy controls (Chaparro et al., 2020; Miranda et al., 2016) whereas the effectiveness and utility of these algorithms should be explored in large-scale studies on various populations with both healthy and pathological patients (Table S1).

Several SP algorithms have been proposed in clinical studies of gait. These algorithms used data from force plates (Song et al., 2020), anatomical markers from motion capture systems (Yoon et al., 2012), marker-less systems (e.g., video, Kinect, ultrasound) (Hunt et al., 2018; Wei et al., 2020). For instance, force plate data enable a controller to calculate spatiotemporal parameters, which can be used to adjust treadmill speed at every time step (Song et al., 2020). Marker-based or marker-less algorithms usually calculate centre of mass (CoM) position as an input to a controller to adjust belt speed. Regardless of the input data, these algorithms aim to adjust the treadmill speed to the subjects’ comfortable gait speed (Bahadori et al., 2021; Canete and Jacobs, 2021; Kimel-Naor et al., 2017; Plotnik et al., 2015).

We present a well-established algorithm for SP treadmills. This algorithm is commercially implemented in the GRAIL, M-Gait and CAREN (Motek Medical B.V., Houten, The Netherlands) and has been studied with different populations (Geijtenbeek et al., 2011). The goal of the algorithm is to keep the subject in the middle of a treadmill while adjusting the treadmill’s speed to match with the subject’s self-selected speed. The specific aims of this paper are: a) to explain an SP controller algorithm for treadmill walking and make it freely available to all and b) to provide an example (MATLAB scripts) of the algorithm that simulates SP walking. We hope this will enable further comparison between various algorithms in the literature.

## Methods

### Algorithm

The self-paced algorithm uses an input 1-D *position* (anterior-posterior or *z* direction) to calculate a treadmill *speed*. The position of the subject is usually determined using the average of four (pelvic) markers, but theoretically a force centre of pressure, or body weight support yoke position could also be used (Yoon et al., 2012).

The algorithm aims to keep the subject near the centre of the treadmill with (*z* ≅ 0), to reach the same subject’s self-selected speed (*v*_*b*_ ≅ *v*_*s*_) and to minimize the ripple effect of the belts *a*_*b*_ → 0. It should be noted that (*v*_*s*_ − *v*_*b*_ = *v*_*com*_ *or ż*). See Table 1 for nomenclature.

**Table 1:** Nomenclature table. The empty cell values are calculated in the experiment using the SP algorithm.

| Parameters | Unit | Parameters | Values |
| --- | --- | --- | --- |
| Position gain | | $g_p$ | 0.3 |
| Speed gain | | $g_v$ | 1.0 |
| Acceleration gain | | $g_a$ | 1.1 |
| Front position | $m$ | $z_f$ | -0.55 |
| Back position | $m$ | $z_b$ | 0.4 |
| Position threshold | $m$ | $T_p$ | 0.03 |
| Filter frequency | $Hz$ | $\omega$ | 1.0 |
| Sensitivity | | $s$ | 1.0 |
| Belt speed limits | $m/s$ | $v_b$ | 0.0 to 5.0 |
| Zero position | $m$ | $z_0$ | |
| Subject's position in AP direction on treadmill | $m$ | $z$ | |
| Subject's filtered position | $m$ | $z_f$ | |
| Subject's filtered speed | $m/s$ | $\dot{z}_f$ | |
| Low-pass filter function | | $f$ | |
| Time (s) | $s$ | $t$ | |
| Distance to the front of treadmill ($m$) | $m$ | $d_f$ | |
| New front position in treadmill relative to subject ($m$) | $m$ | $z_{nf}$ | |
| Distance to the back of treadmill ($m$) | $m$ | $d_b$ | |
| New back position in treadmill relative to subject | $m$ | $z_{nb}$ | |
| Acceleration due to relative position (like P in PD control) | $m/s^2$ | $a_p$ | |
| Acceleration due to relative position (like D in PD control) | $m/s^2$ | $a_s$ | |
| Distance to the front or back | $m$ | $d$ | |
| Subject speed | $m/s$ | $v_s$ | |
| Calculated acceleration in algorithm | $m/s^2$ | $a_{cal}$ | |
| Calculated belt speed in algorithm | $m/s$ | $v_{cal}$ | |

The controller calculates the treadmill speed by incorporating the position and speed of the subject. The treadmill accelerates when the subject is at the front and/or moving to the front of the treadmill and vice or versa. In case the subject is close to the front or back of the treadmill, the speed is quadratically incorporated.

### Configuration

The following configuration parameters are used, with the default values (Table 1). The gains for the three acceleration terms were experimentally adjusted to obtain smooth transition toward a steady state belt speed. The sensitivity values can be adjusted in D-Flow. D-Flow is software that this algorithm is implemented and integrates hardware and software (Geijtenbeek et al., 2011).

### Calculation

At the start of each update, several checks take place e.g.

- Is the last update > 0.5 s?
- Does the input position remain unchanged for > 0.5 s?

Then, given an input position *z* and current time *t*, the following calculations are performed:

First, the input position is filtered using a second order Butterworth filter:

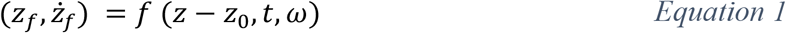

Distances to the back and front of the treadmill are determined and limited (to prevent them from being zero, Equation 2-Equation 3).

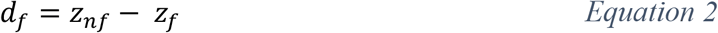

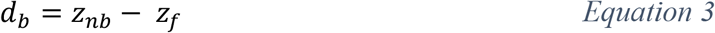

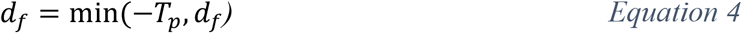

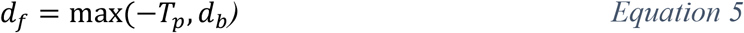

The new speed is based on three acceleration terms (note that *forward* direction is negative, see Figure 1):

**Figure 1:**
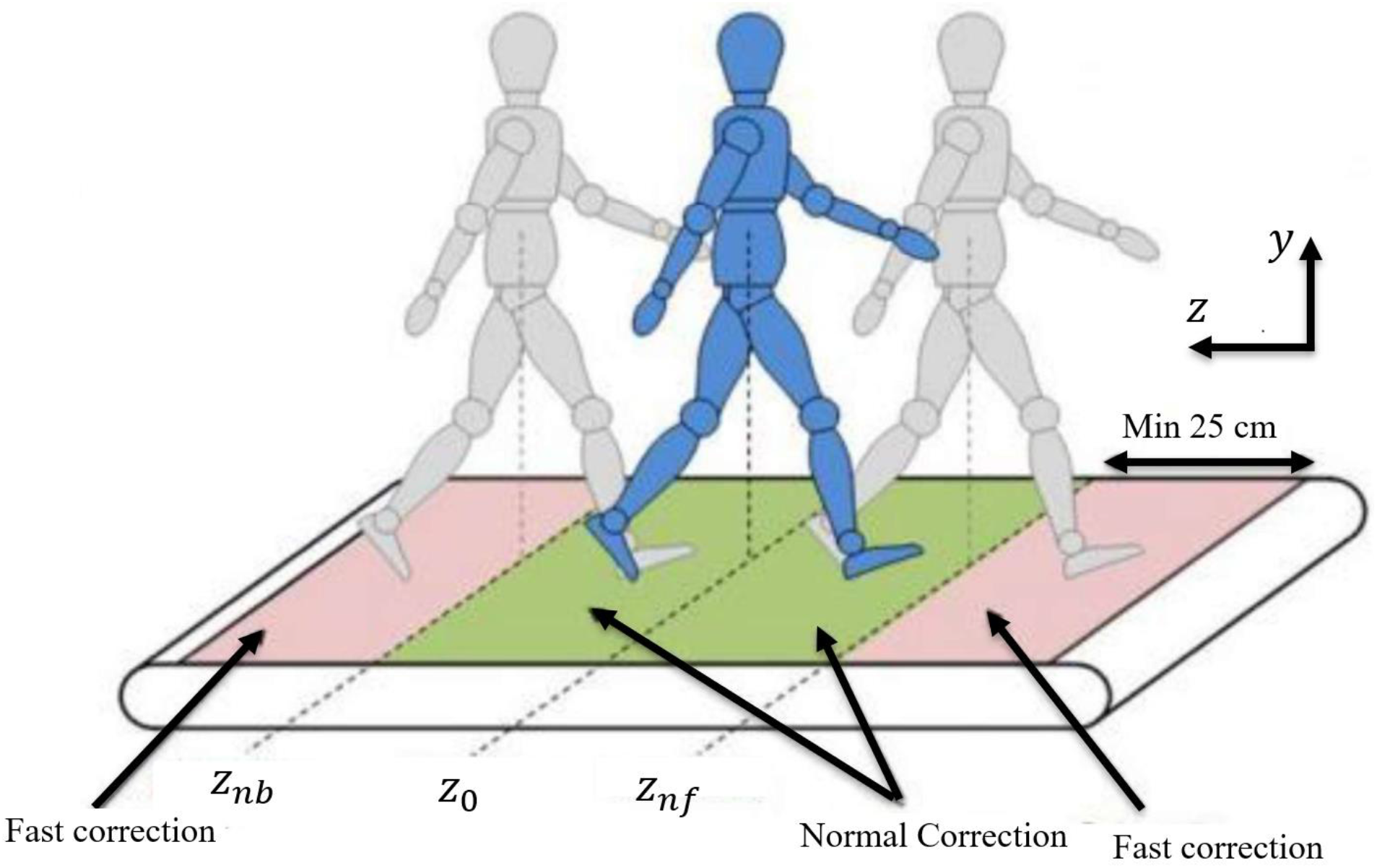
Position of subjects and definition of certain positions explained in the equations. Fast corrections occur as the distance from the front or back (*d*) becomes small thus 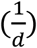 velocity acceleration (*a*_*v*_) increases (see Equation 12)

The *position-based acceleration* is based on the position of the subject: the acceleration increases when the subjects is positioned in the front of the treadmill (Equation 6).

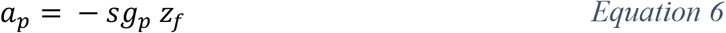

The *speed-based acceleration* is based on the speed of the subject: the acceleration increases when the subject is moving forward. This effect is increased when the subject is further away from the zero position (Equation 7).

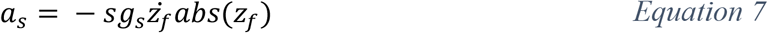

The *velocity-based acceleration* is based on the squared speed of the subject. The effect of this term is increased when the subject is close to the front or back (depending on which direction the subject is moving) of the treadmill. In case the subject is moving forward, and approaches the front of the treadmill, the denominator 2*d* will be small, resulting in an increased acceleration.

If *v*_*s*_ < 0 (i.e., subject is moving forward) then *d* = *d*_*f*_ otherwise, *d* = *d*_*b*_

One can derive the equation below by considering that the subject should be prevented from reaching front and back of the treadmill (*v*_*s*_ → 0) with minimal treadmill acceleration thus 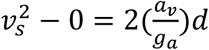 where 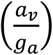 is the minimal acceleration and *g*_*a*_ can be adjusted experimentally, thus:

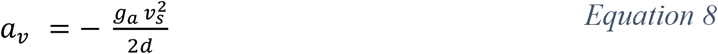

Determine the total acceleration and integrate to arrive at the new speed (Equation 9 & Equation 10)

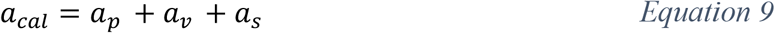

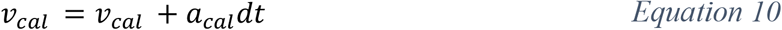

Finally, the calculated speed is limited based on the minimum and maximum speed set by the user (Equation 11).

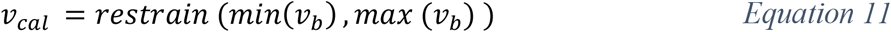

### Simulated example

MATLAB scripts were developed to simulate the algorithm (Supplementary 1). A subject performed a self-paced walking on a GRAIL system (Motek Medical B.V, Houten, Netherlands). Three sensitivities were selected to calculate belt speeds: *s* = 0.2, 1 and 2. The outcome of the function is the belt speed which can be compared to the actual experiment conducted on a treadmill controlled by SP algorithm in D-Flow software. We also calculated the contribution of each acceleration component (Equation 13) during SP algorithm experiment in this specific example.

## Results

The results of belt speed due to three different sensitivities were presented in Figure 2. As the *s* changed, we observed that *s* = 1 presented a pattern where the belt speed reached steady state in about 20s from standing position. Moreover, the belt speed from the experiments matched well with the simulated algorithm in MATLAB. The contribution of each acceleration term to total acceleration of the belt were quite different (Figure 2).

**Figure 2:**
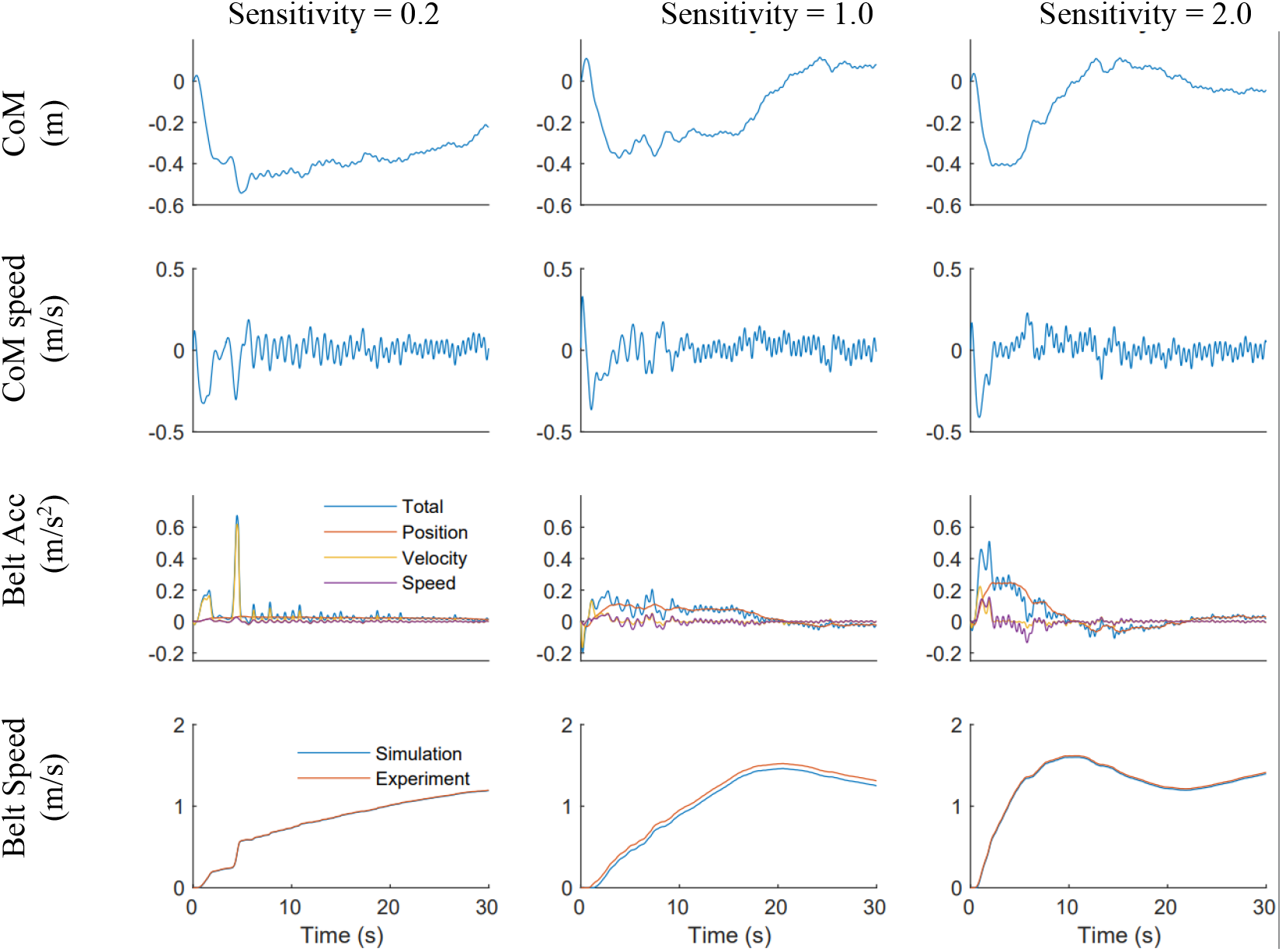
An example of using self-paced algorithm to calculate belt speed based on centre of mass (position and speed) using three different sensitivities. Belt acceleration and contribution of each term (position, velocity and speed, see equation 14), were calculated based on CoM position. Then total belt acceleration is used to calculate belt speed. Although the belt acceleration was not smooth the final belt speed i.e., what subject experiences during self-paced walking are quite smooth.

## Discussion

We presented a self-paced algorithm for an instrumented treadmill that has been tested in a variety of clinical and research settings (Supplementary 2). Using an approximate centre-of-mass position and velocity from motion capture data, this self-paced algorithm calculates belt speed with different sensitivities based on a non-linear controller. Specially three acceleration terms are computed. Belt speed is calculated by integrating sum of these accelerations at each time step. Total belt acceleration was dominated by position-based acceleration when there was a small change in the position on the treadmill.

One key strength of this algorithm is the way belt acceleration is calculated if the subject is close to the front or back of the treadmill (see Equation 15). Based on the subject’s COM data, the algorithm attempts to prevent the subject reaching the extreme front and back positions using minimal treadmill acceleration. As the subject moves toward edges (front or back) of the treadmill, the algorithm brings them back quickly (see Equation 16) and smoothly to the centre of the treadmill. This algorithm has been tested in a number of studies with various populations including healthy, pathological (e.g. CP), old, young, males and females on at least 400 people (Bahadori et al., 2021; Canete and Jacobs, 2021; Kimel-Naor et al., 2017; Plotnik et al., 2015). A key limitation of this algorithm is that it can be used in high performance mode e.g., running, with perturbations, and visual/auditory feedbacks in different populations across the lifespan.

## Future direction

The self-paced algorithm presented in this study can be applied in various areas of human performances including healthy and pathological from numerous disciplines such as rehabilitation, and design of a robotics, exoskeleton system. Table S2 shows many uses of the algorithm (Supplementary 2) indicating its great potential in those areas (Song et al., 2020) and beyond.

## Conclusion

We presented a unique self-paced algorithm based on position of the centre of mass. This self-paced algorithm has been used in various experimental and clinical settings. It has shown a great potential for self-paced studies to closely simulate over ground walking via instrumented treadmill while integrated with Motek devices.

## Supporting information

Supplemental Scripts 1

## Acknowledgement

The authors would like to thank Motek’s development team and special thanks to Craig Leach and Ben van Basten for the early draft of the method section.

## Appendixes

**Supplementary 1:** Scripts that simulate self-paced algorithm in MATLAB (selfPacedTreadmill.m & mox2mat.m) based on an experiment on a GRAIL system (3 text files with hip markers and experimental belt speeds in D-Flow for comparison with MATLAB simulations), and a description of how to run SP in MATLAB and D-Flow (SP_How2RunCodes.docx). These are all in a zip file (7 files including 3 experimental data with different sensitivities plus how to do word document).

**Supplementary 2:** A list of publications using this SP algorithm since 2014.

