## Supplemental Table 2 for "Self-paced treadmill controller algorithm based on position and speed of centre of mass"

**Table S2 (Supplementary 2):** Studies in which the current algorithm has been used from 2014-2022 at the time of writing Feb 2022.

| **# Study** | **Population** | **#Subject** | **Device** |
| --- | --- | --- | --- |
| 1. (Bahadori et al., 2021). | Healthy adults | 100 | GRAIL |
| 2. (Kimel-Naor et al., 2017) | Young healthy adults | 11 | CAREN High-End |
| 3. (Ibala et al., 2019) | Able body healthy (5 female) | 11 | CAREN High-End |
| 4. (Castano and Huang, 2021) | Young adults | 10 | M-Gait |
| 5. (Kao and Pierro, 2021) | Healthy young | 18 | M-Gait |
| 6. (Soangra and Rajagopal, 2021) | Healthy young | 16 | GRAIL |
| 7. (van der Krogt et al., 2014) | Healthy subjects | 19 | GRAIL |
| 8. (Rohafza et al., 2022) | 13 Patients with Parkinson’s Disease and thirteen healthy controls | 26 | GRAIL |
| 9. (Sloot et al., 2015) | 11 typically developing (TD) children and 9 children with cerebral palsy (CP). | 20 | GRAIL |
| 10. (Plotnik et al., 2015) | Young healthy | 26 | V-Gait |
| 11. (Lu and Al-Amri, 2019) | Healthy young | 28 | GRAIL |
| 12. (Theunissen et al., 2022b) (Theunissen et al., 2022a) | Healthy (9 females) | 18 | CAREN |
| 13. (Baron et al., 2018) | Patients with Parkinson's disease | 23 | CAREN |
| 14. (Jeschke et al., 2019) | Healthy young adults | 13 | GRAIL |
| 15. (Van Bladel et al., 2021) | Persons after stroke | 25 | GRAIL |
| 16. (Kimel-Naor et al., 2017) | young healthy | 11 | CAREN High-End |
| 17. (Bugnariu et al., 2013) | 7 pairs of boys with ASD and age-matched controls | 14 | CAREN |
| 18. (Chaparro et al., 2020) | Participants (21 females) | 30 | C-Mill |
| **Total participants** | **Healthy and pathologic** | **419** |  |

Van Bladel, A., De Ridder, R., Palmans, T., Van der Looven, R., Cambier, D., 2021. Comparing spatiotemporal gait parameters between overground walking and self-paced treadmill walking in persons after stroke, in: ESMAC 2021.

van der Krogt, M.M., Sloot, L.H., Harlaar, J., 2014. Overground versus self-paced treadmill walking in a virtual environment in children with cerebral palsy. Gait Posture 40, 587–593.
