## Supplemental Scripts 1 for "Self-paced treadmill controller algorithm based on position and speed of centre of mass": mox2mat.m

```
%% How to read .mox file from D-Flow in Matlab?
% Author: Hossein Mokhtarzadeh
% Date: 23 Feb 2022, v5
% This is a script to convert .mox file from D-Flow to Malab Table (and excel &txt) format
% Thus any manipuation of data inside mox file is possible within Matlab
% Tested in this version (R2021b)!
%% ToDos
% 1. if interest is there, recreate simple pdf report
%% Read the .mox file
function T = mox2mat(filename)

%% Example: 
% filename = 'SF03_sp_slow.mox';T = mox2mat(filename);plot(T.("Belt.Speed"));plot(T.("FP0 - Force Y"))

%% read new converted xml file in Matlab as struct
S = readstruct(filename,'FileType','xml');
%% Create a table 
for i=1:length(S.viewer_data.viewer_channel)
    labls(i) = S.viewer_data.viewer_channel(i).channel_label;
    temp = S.viewer_data.viewer_channel(i).raw_channel_data.channel_data(1,1);
    data(:,i) = cell2mat(textscan(temp, '%f'));
end
T = array2table(data,'VariableNames',labls);

%% Export to excl + txt etc later uncomment if interested
% writetable(T,'sampletable.txt')
% writetable(T,'myData.xlsx');

end
```
