## Supplemental Scripts 1 for "Self-paced treadmill controller algorithm based on position and speed of centre of mass": plotting.m

```
% make three and get what M1 are and just plot all..bearuty this one..
% if more needed some changed needed her...
clc
close all
clear
sensitivity = [0.2 1 2]; % we change them in this paper
files = {'SP_sensitivity0.2.txt';'SP_sensitivity1.txt';'SP_sensitivity2.txt'}; % experiments can also be .mox files record experments from 0 to steady-state
defaults = {-0.1,  -0.55, 0.4, 0.3, 1, 1.1, 0.03, 1, 0.0, 2.0, 300}; % if your treadmill or system has diff parameters, adjust here.
% T1 = readtable(files{1});
% T2 = readtable(files{2});
% T3 = readtable(files{3});
for j=1:3
    [filtered_pos, filtered_speed, y, t, acc, T] = selfPacedTreadmill(files{j},sensitivity(j), defaults); %mox files accepted here
    % title('Belt Speed (m/s)');
    %% First row
    subplot(4,3,j)
    plot(t,filtered_pos);set(gca,'xtick',[]);title('Sensitivity = 0.2');set(gca, 'box', 'off');
    ylim([-0.6 0.2]);
    if j==1
        ylabel('CoM (m)')
        title('Sensitivity = 0.2');
    elseif j==2
        title('Sensitivity = 1');
    else
        title('Sensitivity = 2');
    end

    %% Second row
    subplot(4,3,j+3)
    plot(t, filtered_speed);set(gca,'xtick',[]);set(gca, 'box', 'off');
    if j==1
        ylabel('CoM Speed (m/s)');
    end
    ylim([-0.5 0.5]);

    %% Third row
    subplot(4,3,j+6)
    %     plot(t, T.speed);ylabel('Belt Speed Experiment (m/s)');ylim([0 2]);
    hold on;plot(t, acc(:,1))
    plot(t, acc(:,2))
    plot(t, acc(:,3))
    plot(t, acc(:,4))
    if j==1
        legend({'Total', 'Position', 'Velocity', 'Speed'});legend('boxoff');
        ylabel('Accelerations (m/s^2)');hold off;
    end
    set(gca,'xtick',[]);ylim([-0.25 0.8]);
    % add accel;eraton here and belt speed in another or just compare
    % w/next

    %% Fourth row
    subplot(4,3,j+9)
    hold on;plot(t,y);ylim([0 2]);
    plot(t, T.speed);ylim([0 2]);hold off;
    if j==1
        ylabel('Belt Speed (m/s)')
        legend('Simulation', 'Experiment');legend('boxoff');
    end
    xlabel('Time (s)');
    %     clear filtered_pos filtered_speed y
end

%% save it as high quality image or pdf
% this is great: https://au.mathworks.com/help/matlab/creating_plots/save-figure-at-specific-size-and-resolution.html
set(gcf, 'PaperUnits', 'inches');
set(gcf, 'PaperSize', [6 8]);
f = gcf;
exportgraphics(f,'bartransparent.pdf','ContentType','vector',...
               'BackgroundColor','none')
% see this and make sure all the fonts, size etc r perfect
% https://au.mathworks.com/matlabcentral/answers/593113-exporting-a-figure-including-multiple-subplots-into-high-resolution-pdf-or-svg
% saveas(gcf,'subplots','pdf')
% print('test','-dsvg')
%% acc to b added and velcoty from experimetns

% T1 = readtable(files{1});
% T2 = readtable(files{2});
% T3 = readtable(files{3});
% figure
% subplot(1,3,1)
% plot(T1.Time-T1.Time(1), T1.speed)
% subplot(1,3,2)
% plot(T2.Time-T2.Time(1), T2.speed)
%
% subplot(1,3,3)
% plot(T3.Time-T3.Time(1), T3.speed)

%% Plot accelerations
% figure;hold on
% plot(t, acc(:,1))
% plot(t, acc(:,2))
% plot(t, acc(:,3))
% plot(t, acc(:,4))
% legend({'Total Acc', 'Pos Acc', 'Vel Acc', 'Speed Acc'});
% ylabel('Accelerations (m/s^2)');xlabel('Time (s)')
```
