## Supplemental Scripts 1 for "Self-paced treadmill controller algorithm based on position and speed of centre of mass": selfPacedTreadmill.m

```
%% %            Self-paced treadmill algrithms         %%
%%We present how Motek' Self-Paced algorith works,     %%
%%This uses three terms like PD controller to reach SP %%
%%Steady state as smooth as possible                %%%%%
%%%%%%%%%%%%%%%       12 Mar 2022, v6       %%%%%%%%%%%%%
% Authors: Hossein Mokhtarzadeh, Rosie Richardson, Thomas
%%%%%%%%%%%%%%%%%%%%%%%%%%%%%%%%%%%%%%%%%%%%%%%%%%%%%%%%%


function [filtered_pos, filtered_speed, y, t, acc, T] = selfPacedTreadmill(filename, sensitivity, defaults)

    %% ToDo:
    % table done what else?
    % 1. clean the final figures
%     change dialog since we have 3 s so easy to plot now
    % 2. any other figures we aim to put in the paper?
    % 3. the reason for input pop up is that marker names are diff in diff files.
    % if not it is easier to just find pelvic markers

    %% Example
    % see plotting.m for more info, filename can be .mox or .txt
    % filename = 'Trial_0002.txt'; see mox2mat.m if mox is the input

    %% Parameters
    % defaults = {-0.1,  -0.55, 0.4, 0.3, 1, 1.1, 0.03, 1, 0.0, 2.0, 300};
    [zero_pos, new_front_pos, new_back_pos, new_pos_gain, new_speed_gain,...
        new_acceleration_gain, new_pos_threshold, new_filter_frequency, ...
        min_speed, max_speed, sampling_rate] = deal(defaults{:});
    arbitaryDis = 0.0; % change the data and pos to move toward front or end of forceplates, this is arbiraty, it must be Zero in fact.!
    calculated_speed = 0;


    %% 0. read datac can b table and header read later

    [~,~,ext] = fileparts(filename);
    if strcmp(ext, '.mox')
        T = mox2mat(filename);
    else
        T = readtable(filename);
    end


    %% if you want to select variables, use this below
%     list = T.Properties.VariableNames;
%     [indx,~] = listdlg('PromptString',{'Select pelvis markers in AP direction.',...
%         'Four markers in Z direction & Use Ctr to select all.',''},...
%         'ListString',list);
%     pos = (T.(indx(1)) + T.(indx(2)) + T.(indx(3)) + T.(indx(4)))/4;
% 
%     [indx,~] = listdlg('PromptString',{'Select time column.'},...
%         'SelectionMode','single', 'ListString',list);
% 
%     t = T.(indx);
%     t = t-t(1); % start from t0

    %% since we have new data from Motek GRAIL system with following columens: Time	speed	M1z	M2z	M3z	M4z so simplify things for plots
    pos = (T.M1z + T.M2z + T.M3z + T.M4z)/4;
    t = T.Time;
    t = t-t(1); % start from t0

    %% 1. Low pass filter second order. make changes here if other filters needed!
    F = new_filter_frequency;
    Fs = sampling_rate;
    [y, x] = butter(2, F/(Fs/2));

    %% start algorith
    filtered_pos = filter(y,x, pos-zero_pos);filtered_speed = gradient(filtered_pos)./gradient(t);

    %% 2. distance_to_front = new_front_pos - filtered_pos
    % forward direction (-)

    distance_to_front = new_front_pos - filtered_pos; % plot(distance_to_front)
    distance_to_back  = new_back_pos - filtered_pos;% plot(distance_to_back)
    distance_to_front = min(-new_pos_threshold, distance_to_front);
    distance_to_back = max(new_pos_threshold, distance_to_back);

    %% 3. New speed is current speed + sum of acceleration terms:
    pos_acc = sensitivity * new_pos_gain * (-filtered_pos);
    speed_acc = sensitivity * new_speed_gain * (-filtered_speed) .* abs(filtered_pos);

    d=zeros(length(filtered_speed),1);
    for i = 1:length(filtered_speed)
        if filtered_speed(i) < 0 % i.e. (subject is moving forward)
            d(i) = distance_to_front(i);
        else
            d(i) = distance_to_back(i);
        end
    end

    %% 4. Calculate belt speed

    vel_acc = new_acceleration_gain * (filtered_speed .^2) ./ (2 * (-d));
    calculated_speed = 0;
    calculated_acc = pos_acc + vel_acc + speed_acc;

    for i = 2:length(filtered_speed) % just starting from 2 cos of derivation or i-1 in fact!
        calculated_speed(i) = calculated_speed(i-1) + (calculated_acc(i) * (t(i)-t(i-1)));
    end

acc = [calculated_acc pos_acc  vel_acc speed_acc];

    %% 5. Finally, the calculated speed is limited based on the minimum and maximum speed set by the user.
    y = bound(calculated_speed, min_speed, max_speed);

function y = bound(x,bl,bu)
    %return bounded value clipped between bl and bu
    y=min(max(x,bl),bu);
```
