## Supplemental Scripts 1 for "Self-paced treadmill controller algorithm based on position and speed of centre of mass": SP_How2RunCodes.docx

**Supplementary 1:**

**How to run the codes in this paper?**

We provide two sets of codes: one in MATLAB and the other one in D-Flow. Since using D-Flow requires additional license we provided the MATLAB files so that everyone can run and use the codes as they wish.

1. **MATLAB code**

You are given 6 files including an experimental data (3 files with different sensitivities *SP_sensitivity*.txt*) to reproduce the results in this paper MATLAB.

If you run the plotting.m the results of this paper will be recreated.

a) *selfPacedTreadmill.m*

This MATLAB file needs input of treadmill information (sensitivity and defaults) and the .mox or .txt files where marker data is available. Mox files are xml format files implemented in Motek devices. One can think of equivalent of c3d files where data is stored in different channels.

[filtered_pos, filtered_speed, y, t] = selfPacedTreadmill(sensitivity, defaults)

After running this, the user needs to select the pelvis marker (4 of them) so that the algorithm calculates belt speed.

b) *mox2mat.m*

This file can be standalone or be an input the script above. If the input file is in the .mox file, this code provides a table readable by MATLAB from the data available in the .mox file.

*Note: Make sure all the marker data are in this .mox file so that the codes work properly.*

c) *plotting.m*

This file recreated all the figures in this paper.

*How to use and run these codes?*

You can simply copy all these files to a folder you wish! Then you can run them easily. The codes were tested in (R2021b) Update 2.

2. **D-Flow** (software and if experimental hardware of Motek needed so not free)

To activate the safe-paced mode in Motek devices the following program can be used. MoCap module can streamline the marker data. Expression module averages the pelvic markers’ position in z direction i.e., Anterior/Posterior direction. This average value will be input to the treadmill for the SP mode as shown below. Users can change the sensitivity of the controller in the GUI.

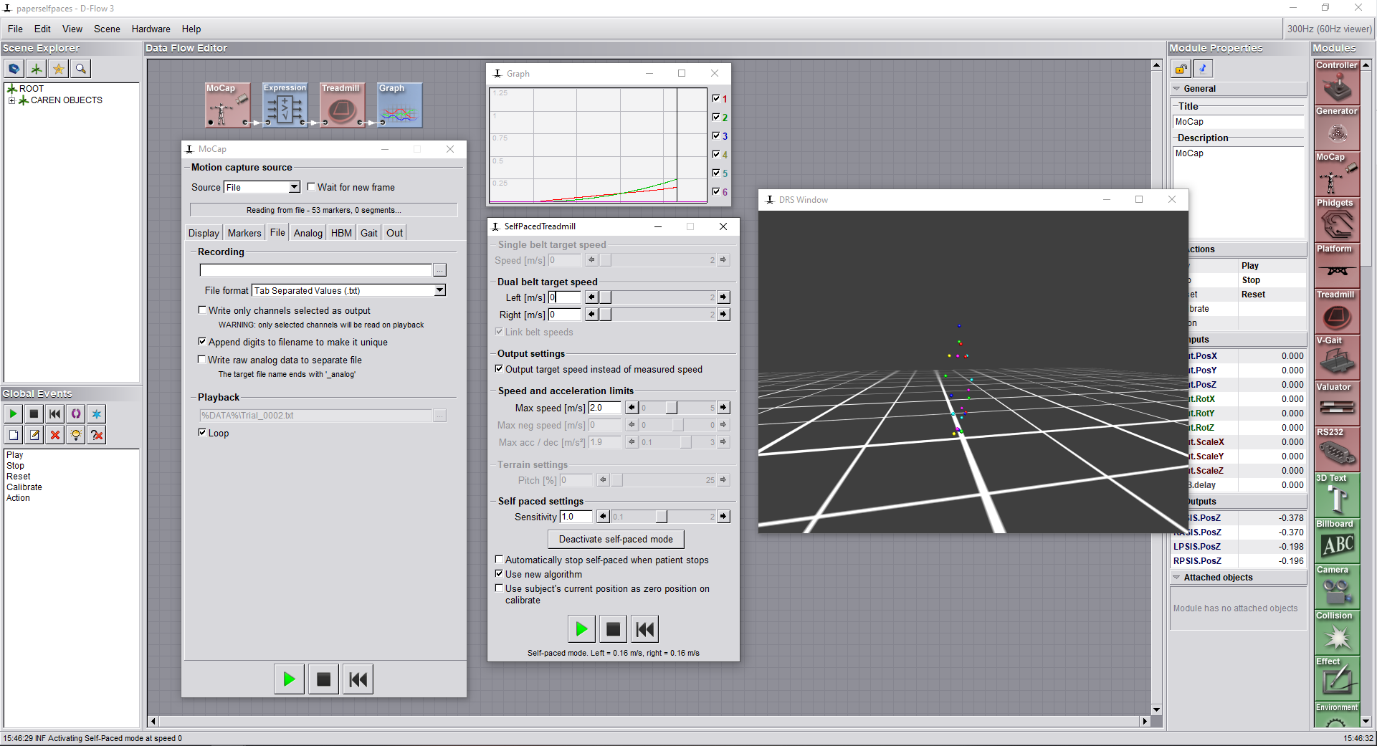

Figure 1: how to run and activate SP in D-Flow with an example. The package in this paper can be used and unpacked in D-Flow to reproduce the outcomes here.

Also, we created a short video to present how the self-paced algorithm is implemented and performed in D-Flow: <https://tinyurl.com/MotekSelfPaced>
